# Consistent variations in personality traits and their potential for genetic improvement of biocontrol agents: *Trichogramma evanescens* as a case study

**DOI:** 10.1101/2020.08.21.257881

**Authors:** Lartigue Silène, Yalaoui Myriam, Belliard Jean, Caravel Claire, Jeandroz Louise, Groussier Géraldine, Calcagno Vincent, Louâpre Philippe, Dechaume-Moncharmont François-Xavier, Malausa Thibaut, Moreau Jérôme

## Abstract

Improvements in the biological control of agricultural pests require improvements in the phenotyping methods used by practitioners to select efficient biological control agent (BCA) populations in industrial rearing or field conditions. Consistent inter-individual variations in behaviour (i.e. animal personality) probably affect BCA efficiency, but have never been taken into account in the development of phenotyping methods, despite having characteristics useful for phenotyping: repeatable (by definition), often heritable, etc. We developed a video-tracking method targeting animal personality traits and evaluated the feasibility of its use for genetic improvement in the BCA *Trichogramma evanescens*, by phenotyping 1,049 individuals from 24 isogenic lines. We found consistent individual variations in boldness, activity and exploration. Personality differences between the 24 isogenic lines suggested a genetic origin of the variations in activity and exploration (broad-sense heritability estimates of 0.06 to 0.11) and revealed a trade-off between exploration and fecundity.

## Introduction

The demand for more sustainable agriculture is increasing worldwide (Godfray et al., 2010; Willer & Lernoud, 2019). Various elements can be used in the development of sustainable strategies, and biological control (BC) is one such element that is currently attracting considerable attention (van Lenteren 2012). Most BC methods are based on the choice, rearing and introduction of biological control agent (BCA) populations able to control the target pests (Eilenberg, Hajek, & Lomer, 2001). Choosing the right BCA is key to the success of pest regulation programmes and is based on (i) the ability of the BCA to control pest populations in the field, (ii) its potential to adapt to the release environment, (iii) its expected impact on local biodiversity, and (iv) the feasibility of mass-rearing and storing the BCA in industrial conditions (Briese, 2000; Kruitwagen, Beukeboom, & Wertheim, 2018; Sforza, 2010). The identification of BCA species or populations with as many of the desired features as possible is time-consuming and complex, particularly given that the choice of non-indigenous species before use as BCAs is constrained by increasingly strict regulations for the protection of biodiversity (Lommen, Jong, & Pannebakker, 2017).

Phenotyping is key for (i) the efficient characterisation of traits related to the desirable features of BCAs listed above, (ii) smart choices of BC taxa when screening the available natural enemy diversity and (iii) the management of phenotypic evolution in industrial contexts involving rearing procedures and quality control (Kruitwagen et al., 2018; Lommen et al., 2017). However, the phenotyping methods currently used in the choice of BCAs or for quality control are mostly low-throughput and based on single proxies of fitness, such as predation or parasitism rate, size, sex ratio, longevity, or developmental rate (Hopper, Roush, & Powell, 1993; Prezotti, Parra, Vencovsky, Coelho, & Cruz, 2004; Roitberg, Boivin, & Vet, 2001; Smith, 1996). These proxies are intuitively correlated with fitness under laboratory conditions, but their actual relevance for biocontrol, in industrial mass-rearing or field conditions, remains a matter of debate (Lommen et al., 2017; Roitberg et al., 2001). This situation calls for drastic improvements in the phenotyping capacities of the community involved in BC research and innovation.

Behavioural traits are among the most promising of the traits to which more attention could be paid in BCA phenotyping procedures. Most behavioural traits are likely to affect the performance of BCA both during industrial mass rearing and in the field (Roitberg, 2007; Wajnberg, 2009; Wajnberg, Roitberg, & Boivin, 2016). Indeed, studies of BCA behavioural traits have suggested that these traits could (i) facilitate the selection of BCAs that are specific to the targeted pest, (ii) improve release strategies (through studies of the BCA response to pre-release handling or BCA mating behaviour, for example), and (iii) predict the efficiency of target pest suppression by the BCA (Mills & Kean, 2010). However, there have been few studies of BCA behavioural traits for the development of phenotyping methods, and behaviour has been largely neglected by those using BC (Wajnberg, Bernstein, & Alphen, 2008).

As a consequence, the current state-of-the-art for insect behavioural studies displays several key limitations. The first limitation is the lack of diversity of possible target traits for phenotyping. Indeed, although many studies have focused on traits relating to foraging behaviour (Lirakis & Magalhães, 2019; Mills & Wajnberg, 2008), tools for measuring other aspects of behaviour remain scarce. A second limitation is the insufficient focus on the intraspecific variation of traits. Such variation has been comprehensively investigated for only a limited number of BCA species and a limited number of traits (Kruitwagen et al. 2018; Lirakis and Magalhães 2019) (but see however Dumont, Aubry, & Lucas, 2018; Dumont, Réale, & Lucas, 2017; Nachappa, Margolies, Nechols, & Campbell, 2011; Nachappa, Margolies, Nechols, & Morgan, 2010). This situation is detrimental because the investigation of only a fraction of the available intraspecific variability makes it difficult to identify the populations displaying the highest performance for biocontrol, and prevents the development of efficient genetic improvement programmes based on selective breeding and controlled evolution (Wajnberg 2004; Bolnick et al. 2011; Lommen et al. 2017; Kruitwagen et al. 2018, Lirakis and Magalhães 2019). A third limitation is the reliance of most choices in BC exclusively on comparisons between average trait values for species or populations (Lommen et al., 2017). Published studies have suggested that individual variation can affect the characteristics of the population thought to be important for BC (Biro & Stamps, 2008; Michalko, Pekár, & Entling, 2019; Réale, Reader, Sol, McDougall, & Dingemanse, 2007; Wolf & Weissing, 2012).

One way to overcome each of these three limitations would be to apply approaches used in the field of animal personality to BC. Indeed, these approaches provide a framework offering (i) sets of behavioural traits rarely studied in BC and displaying features (repeatability, heritability) that make them good candidates for use in genetic improvement for BC, and (ii) phenotyping methods suitable for analyses of intraspecific variation, including inter-individual variation. Animal personality research focuses on inter-individual differences in behaviour that are consistent over time and context (Dingemanse, Kazem, Reale, & Wright, 2009; Denis Réale et al., 2007). Interest in animal personality has increased over the last few decades, and studies have been performed on diverse taxa, including insects (Amat, Desouhant, Gomes, Moreau, & Monceau, 2018; Bell, Hankison, & Laskowski, 2009; Dingemanse et al., 2009; Gosling, 2001; Kralj-fiser & Schuett, 2014; Mazué, Dechaume-Moncharmont, & Godin, 2015; Monceau et al., 2017; Denis Réale et al., 2007; Sih, Bell, & Johnson, 2004; van Ooers & Sinn, 2011) and, more specifically, insects used as BC agents (Gomes, Desouhant, & Amat, 2019; Michalko et al., 2019). Réale et al. (2007) described five main categories of personality traits: boldness, exploration, activity, aggressiveness and sociability. Boldness represents an individual’s reaction to a risky but not new situation. Exploration is defined as an individual’s reaction to a new situation. Activity reflects the general level of activity of an individual. Finally, in a social context, aggressiveness corresponds to an individual’s agonistic reaction to his conspecifics, and sociability provides information on an individual’s reaction to the presence or absence of con-specifics. Personality traits have been shown to be correlated with traits relevant for pest control, such as foraging capacity, fecundity, growth, survival (Biro & Stamps, 2008), dispersal ability (Cote, Fogarty, Weinersmith, Brodin, & Sih, 2010) and insecticide resistance (Royauté, Buddle, & Vincent, 2014). These traits are probably, therefore, of interest in the context of BC. Moreover, personality traits are repeatable, by definition, and can be heritable (Dochtermann, Schwab, & Sih, 2014; Denis Réale et al., 2007; Stirling, Reale, & Roff, 2002), making them suitable tools for genetic improvement. From a methodological point of view, animal personality provides valuable information for the design of phenotyping and genetic improvement strategies in BC. Indeed, animal personality studies are based on standardised methods designed to measure inter-individual variation and to investigate correlations between traits (e.g. by looking for behavioural syndromes) (Denis Réale et al., 2007; Sih et al., 2004). This is particularly relevant to the objective of selecting several combined BC traits rather than a single trait, as recently recommended by Lommen et al. (2017) and Kruitwagen et al. (2018). The investigation of correlations between traits is also important, to detect trade-offs that may constrain genetic improvement programmes or affect BC traits if mass-rearing causes uncontrolled trait selection (Mackauer, 1976).

In this study, we assessed the potential for BCA phenotyping based on concepts and methods used in the field of animal personality. We used the egg parasitoid *Trichogramma evanescens* Westwood, 1833 (Hymenoptera: Trichogrammatidae) as a test species. *Trichogramma* micro-wasps are used worldwide in augmentative BC against lepidopteran pests (Hassan, 1993; van Lenteren, 2012). Their economic importance (Robin & Marchand, 2020; Thibierge, 2015) justifies investments in research and development aiming to improve their genetic potential. Our aims were (i) to determine whether behavioural traits meeting the criteria of personality traits could be measured in these micro-wasps of approximately 0.5 mm in length; (ii) to investigate the relationships between personality traits and traits classically measured on BCAs in the laboratory, and (iii) to determine whether personality traits could be used in genetic improvement strategies for BCAs. We developed a method based on the video-tracking and measuring, at individual level, of multidimensional behavioural traits relating to boldness, activity and exploration. We investigated the relationship between these behavioural traits and further tested whether these traits were related to individual fitness traits relevant to mass rearing (offspring number, longevity, tibia length). We then compared the traits between 24 near-isogenic strains, to obtain a first insight into the broad-sense heritability of these traits. We looked for genetic correlations potentially constraining the use of these traits for genetic improvement.

## Methods

### Laboratory rearing of T. evanescens

All applicable international, national, and/or institutional guidelines for the care and use of animals were followed. We used 24 near-isogenic lines (hereafter referred to as “lines”) of *Trichogramma evanescens*. Lines were created from inbred crosses in populations established from individuals sampled in different parts of France (geographic origins detailed in Table 7 in the appendix), from 2010 to 2016, and reared in the laboratory at 18 ± 1 °C, 70 ± 10% RH and 16:8 h L:D (details of the protocol followed to create the lines are provided in the appendix). Genetic diversity within lines was below 1.1 alleles per locus at 19 microsatellite loci (unpublished data), and individuals within lines were considered genetically identical. We created two sublines for each line (Lynch & Walsh, 1998), to disentangle the confounding effects of rearing tubes and lines (which may be caused by maternal effects). We considered variation between lines to be of genetic origin, and variation within lines to be of environmental origin. We reared *Trichogramma evanescens* individuals on sterilised *Ephestia kuehniella* Zeller 1879 (Lepidoptera: Pyralidae) eggs, renewed every 10 days, at 25.5 ± 1 °C, 70 ± 10% RH and 16:8 h L:D (Schöller & Hassan, 2001). We kept populations in glass tubes (height: 73 mm, diameter: 11 mm), and fed adults with honey *ad libitum*.

**Table 7.**
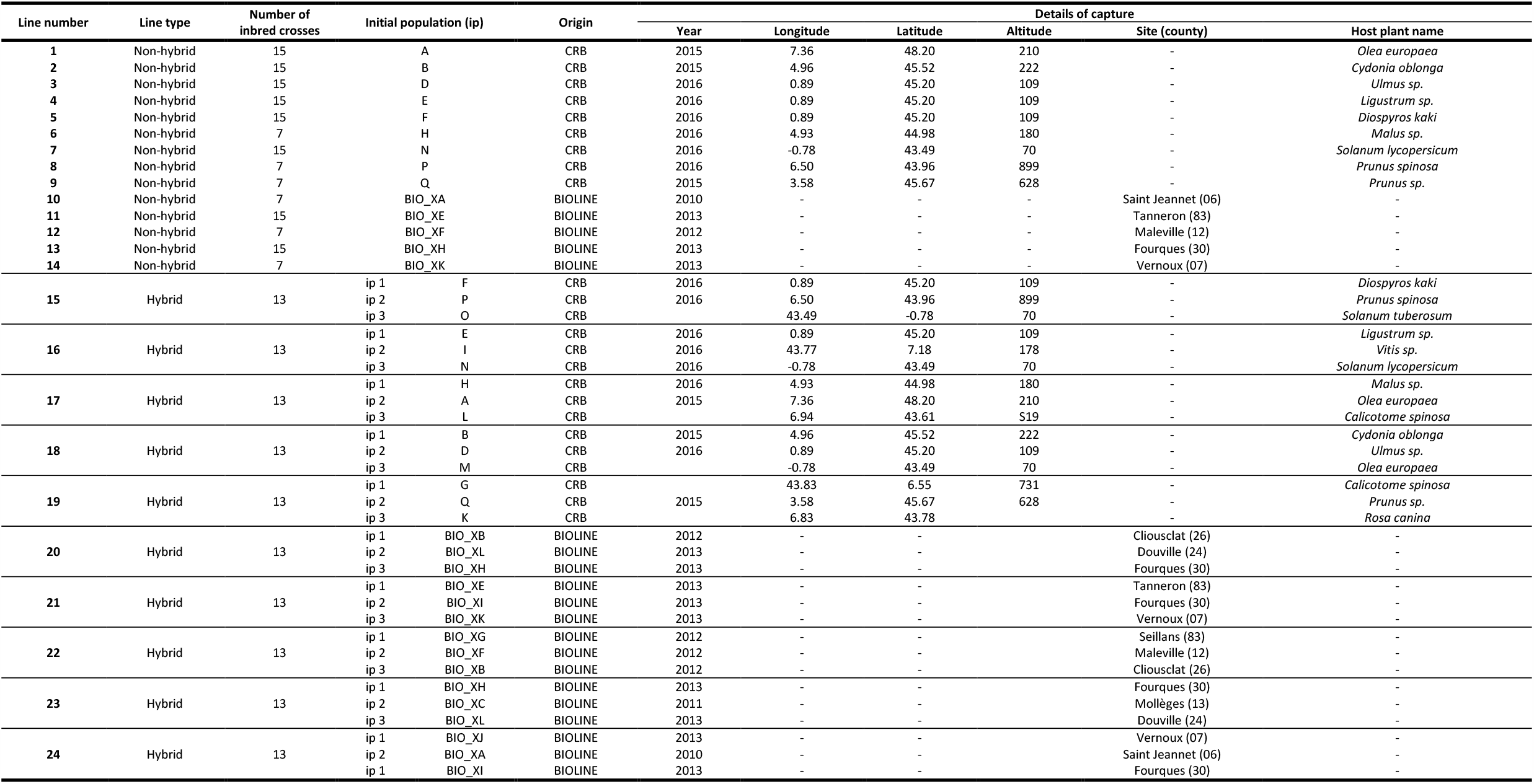
Details on the origins and creation of the lines. Individuals were sampled by the INRAE “Egg-Parasitoids Collection” (CRB EP-Coll, Sophia Antipolis) biological resource centre or the company Bioline AgroSciences Ltd.

### Measurement of variables

#### General experimental design

The following experimental design was used to measure phenotypic traits in *Trichogramma* females (Figure 1). We used mated *T. evanescens* females that had emerged within the last 24 hours, randomly chosen from each line. We checked the physical integrity of these females, which were isolated in glass tubes before the beginning of the experiment (height: 73 mm, diameter: 11 mm) and fed with honey, *ad libitum*. On the first two days, we assessed the behavioural traits of the females. We estimated the number of offspring on days 3 to 5, and longevity from day 6. The experiment lasted from May to July 2019 (about six generations of *T. evanescens*), and was split into 17 experimental sessions, in each of which, we used three females per line. The physiological, developmental and behavioural traits of *Trichogramma* wasps, and of *T. evanescens* in particular, are dependent on temperature (Ayvaz, A., Karasu, E., Karabörklü, S., Tunçbilek, 2008; Schöller & Hassan, 2001). Moreover, as Suverkropp et al. (2001) showed that *T. brassicae* individuals have similar levels of activity throughout the day at temperatures of about 25 °C or higher, we assumed that our *T. evanescens* individuals had similar responses to temperature throughout the day. Therefore, we performed the behavioural experiments at 25.5 ± 1 °C, 70 ± 10% RH. We then measured female longevity and offspring number at 18 ± 1 °C, 70 ± 10% RH, to ensure that the females would live long enough for the final stages of the experiment (Cônsoli & Parra, 1995; Schöller & Hassan, 2001).

**Figure 1.**
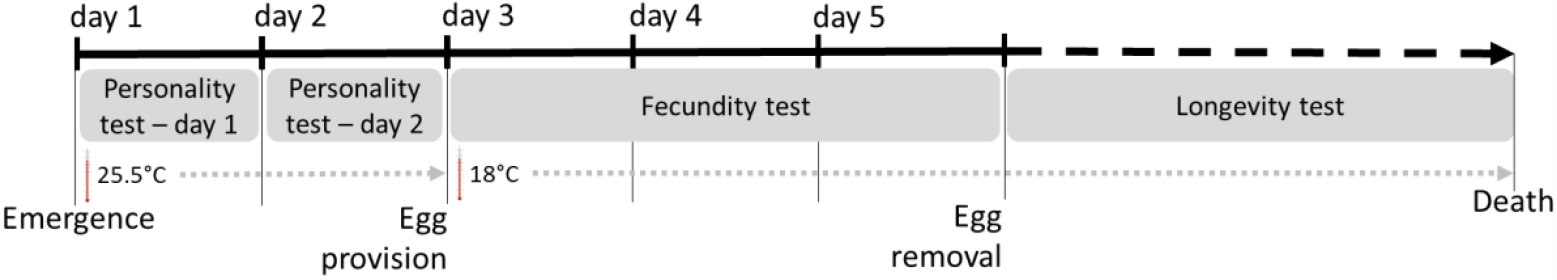
Overview of the experimental design, for one session.

#### Behavioural trait measurement

We observed individuals in an arena composed of two sheets of glass (24 cm × 18 cm), one for the floor and one for the ceiling. The 2 mm space between them was maintained by walls made of a black rubber sheet. We placed this arena on an LCD screen (Samsung© 28” LED, 3840*2160 pixels, 60 Hz) which was used to display a white circle with a diameter of 5.5 cm on a dark background (Figure 2.a). The LCD screen was turned on one hour before the beginning of the experiment, to ensure that a stable temperature of 25.5 ± 1 °C was achieved in the area. The conditions in the growth chamber in which the experimental design was set up were as follows: 22.5 ± 1 °C and 70 ± 10% RH. We used a fine paintbrush to introduce a randomly chosen female into the centre of the arena while the screen was showing a white background. The glass ceiling was replaced, and we then switched to a background with a white circle on a dark background, with the female positioned in the middle of the white circle. We observed the behaviour of the female for 90 seconds, with video recording at 25 frames per second (with a resolution of 1080 p), with a Nikon^©^ D750 camera (Figure 2.a).

**Figure 2.**
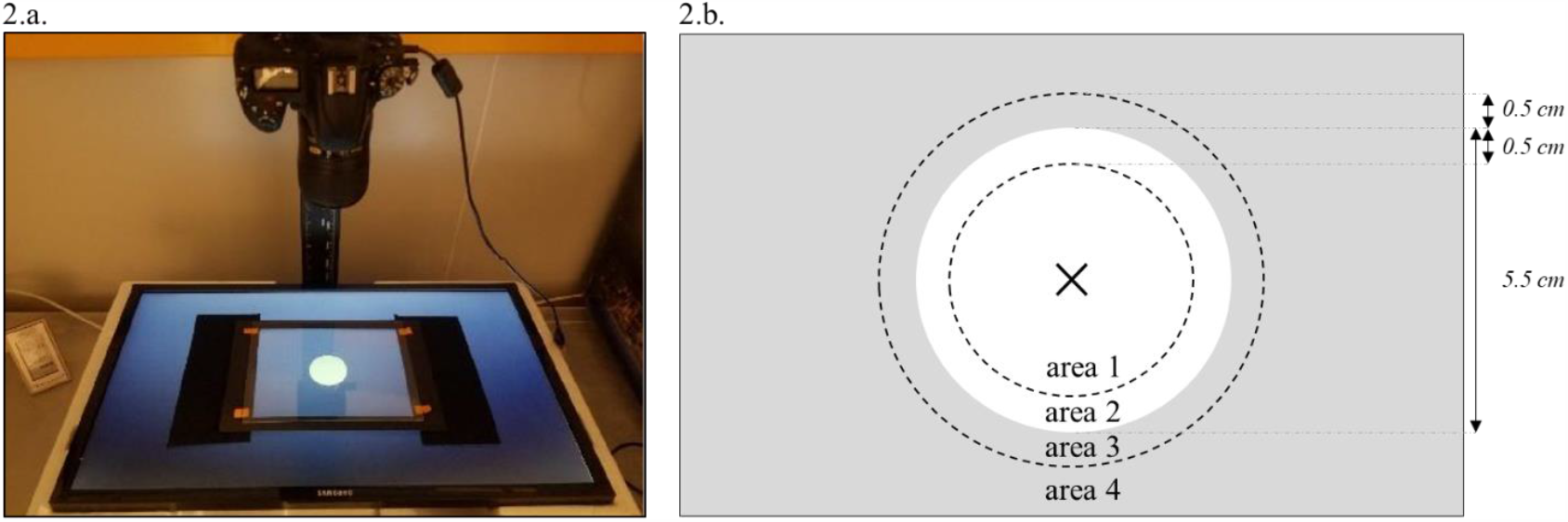
Experimental set-up of the behavioural experiment. Fig. 2.a. shows a photograph of the experimental setup: the LCD screen displaying the white circle on a dark background, the arena and the Nikon© D750 camera above. Fig. 2.b. represents the defined areas of the arena. The grey shading corresponds to the dark background, the white part indicates the white circle, and the dark cross is the site at which the female was placed at the start of the experiment. The dotted lines represent the virtual borders defined between areas 1 and 2, and between areas 3 and 4. The three variables we measured to estimate boldness were (i) the first time until the female first entered the dark area (area 3), (ii) the absolute difference in speed between areas 2 and 1, and (iii) the absolute difference in sinuosity between areas 2 and 1. Both variables we used to estimate exploration (the total area explored per unit time and the mean sinuosity of the pathway) were measured in area 1. Finally, both variables we used to estimate activity (the proportion of time the female was considered to be active and the mean speed) were measured in area 4, so exploration and activity were measured in different areas of the experimental arena.

We then analysed the videos files, determining the 2D spatial position (*x-y* coordinates) and body orientation (in radians) of the female on each frame, with C-trax software (Branson, Robie, Bender, Perona, & Dickinson, 2009). We independently determined the exact position of the border between the white circle and the black background with ImageJ software (Abràmoff, Magalhães, & Ram, 2004). We thus defined regions of interest of 0.5 cm on either side of the border, for investigation of the behaviour of the insect near the border (Figure 2.b). We imported the C-trax and ImageJ output files into R software v.3.6.1 (R Core Team 2019) and cleaned our data to remove tracking artefacts. We used the “trajr” package (Mclean & Volponi, 2018) to calculate speed and sinuosity, in each region of interest. We calculated seven variables we considered to be linked to three personality traits — boldness, exploration and activity — as defined by Réale et al. (2007). As we measured all the variables linked to the three personality traits in the same arena (for feasibility reasons, considering the lifespan of individuals in particular), we decided to measure each variable set linked to each personality trait on a different area of the arena to increase their extent of independence.

Boldness is the reaction of the individual to a risky situation (Réale et al., 2007). We estimated boldness by measuring three variables. The first was the time until the female first entered the dark area (area 3 in Figure 2.b). Higher values indicated that the female took longer to cross the border, which we interpreted as meaning that the female was less bold. The second and third variables were the absolute difference in speed between areas 2 and 1 (Figure 2.b) and the absolute difference in sinuosity between areas 2 and 1 (Figure 2.b). We considered high values for these two variables to indicate a larger change in behaviour at the border, which we interpreted as meaning that the female was more affected by the border and was, therefore, less bold.

Exploration represents the individual’s reaction to a new environment (Réale et al., 2007). Exploration was estimated in area 1 as (1) the total area explored per unit time, and (2) the mean sinuosity of the pathway (Figure 2.b). For this variable, we hypothesised that the females with the most winding pathways remained closer to their release point, indicating a lower level of exploration.

Finally, we measured activity in area 4. Activity was estimated as (i) the proportion of time the female was considered to be active (with a speed of more than 0.01 centimetres per second), referred to hereafter as “activity rate”, and (ii) mean speed (Figure 2.b), considering faster movement to be indicative of a higher level of activity.

We estimated the repeatability of measurements, by conducting two observations per female, with 24 hours between the two measurements, a time interval corresponding to 20% of the mean lifespan of this species. Females were tested in a random order on day 1, and then in the same order on day 2, to ensure that all individuals had exactly the same time interval between two measurements. Between behavioural experiments, each female was placed back in its glass tube and fed with honey, *ad libitum*, in a growth chamber at 25.5 ± 1 °C, 70 ± 10% RH and 16:8 h L:D. Behavioural trait measurements were obtained for 776 to 996 females in total from the 24 lines.

#### Offspring number, longevity and tibia length measurement

After the second day of behavioural observation, females were kept in their glass tubes at 18 ± 1 °C, 70 ± 10% RH and 16:8 h L:D and fed with honey, *ad libidum*. We provided each female with a piece of paper 4.50 cm × 0.85 cm in size, covered with *E. kuehniella* eggs, *ad libidum. E. kuehniella* eggs were removed 72 hours later and placed in conditions of 25.5 ± 1 °C, 70 ± 10% RH and 16:8 h L:D. Three days later, we counted the number of parasitised eggs (considered as black eggs), to estimate the size of the progeny of each female over a period of 72 hours, providing a proxy for female fitness. From day 6, we measured female longevity (the females were still kept in the same individual tubes with honey, but with no *E. kuehniella* eggs, at 18 ± 1 °C, 70 ± 10% RH and 16:8 h L:D). Tubes were checked every day at 5 p.m., until the death of the female. Dead females were conserved in ethanol, for subsequent measurement of tibia length on a micrograph (obtained with an Axioplan II microscope), with ImageJ software (Abràmoff et al., 2004). Images were acquired at the Microscopy Platform of Sophia Agrobiotech Institute, INRA, UNS, CNRS, UMR 1355-7254, INRA PACA, Sophia Antipolis. Not all individuals lived long enough for all the phenotypic measurements to be made. We therefore collected progeny measurements for 929 females, longevity measurements for 655 females and tibia size measurements for 959 females, from all 24 lines.

### Data analysis

We used the R software v.3.6.1 for all statistical analyses. For each variable, we first fitted a linear mixed model with the lme4 package (Bates, Maechler, Bolker, & Walker, 2015), with individual, line, subline and session as random effects. For each variable, data transformations were chosen after graphical inspection of the distribution of model residuals, estimated with the “simulateResiduals” function of the DHARMa package (Hartig, 2019). We performed logarithmic transformations for all behavioural variables except for the area explored within area 1. We addressed several questions regarding the data, and the data analysis for each of these questions is presented below.

#### Are the measured behavioural traits repeatable?

We first estimated the repeatability of the behavioural traits measured with generalised linear mixed models, using the rptR package (Stoffel, Nakagawa, & Schielzeth, 2017). The “rptGaussian” function of the rptR package was used to provide repeatability estimates. As repeatability can be defined as the proportion of variation explained by between-individual variation (Nakagawa & Schielzeth, 2010), we included only two random effects in these models: individual (assuming that the effects of line and subline on variation were included in the individual effect) and session, with individual as a grouping factor. In subsequent analyses, we considered only traits that were significantly repeatable.

#### Do the measured traits identify individual behavioural strategies?

Based on methods generally used in animal personality studies, we first investigated correlations between behavioural traits and then summarized the data by principal component analysis (PCA). We first obtained a single value for each trait for each individual, by extracting, from the linear mixed model described above, linear predictors for each individual, with the “ranef” function of the lme4 package. We used these values to measure the phenotypic correlation between traits, by calculating Spearman’s rank correlation coefficients, to determine whether individuals adopted different strategies, or whether it was possible to describe behavioural syndromes. We estimated bootstrapped 95% confidence intervals from 1000 bootstraps, to assess the significance of the Spearman’s rank correlation coefficients obtained (Nakagawa & Cuthill, 2007), using the “spearman.ci” function of the RVAideMemoire package (Hervé, 2020). *P*-values were adjusted by the false discovery rate method (Benjamini & Hochberg, 1995). We then performed PCA with the “PCA” function of the FactoMineR package (Le, Josse, & Husson, 2008), using both values obtained for each individual (days 1 and 2, when available). We estimated two synthetic personality scores based on the first two axes of the PCA. We used the “fviz_pca_biplot” function of the factoextra package (Kassambara & Mundt, 2019) to obtain a graphical representation of the correlation between repeatable behavioural traits and the distribution of individual values along the two first axes of the PCA.

#### Are the measured traits correlated with fitness-related traits?

We studied the correlation between behavioural and fitness-related traits, using the same linear mixed model as described in the introduction to this section. We extracted linear predictors (using the “ranef” function of the lme4 package (Bates et al., 2015)) for each individual and each personality score from this model. We assessed the correlation between the linear predictors of these personality traits and scores, and offspring number, body size and longevity, by calculating Spearman’s rank correlation coefficients. We estimated bootstrapped 95% confidence intervals to assess significance of the Spearman’s rank correlation coefficients obtained, with the same R function and method as described above. *P*-values were adjusted by the false discovery rate method.

#### Are the measured traits heritable?

We sought to establish a first estimate of broad-sense heritability for each trait. To this end, we followed the simple design proposed by Lynch and Walsh (1998) for clonal populations, and approximated the proportion of the variance explained by genetic factors with an estimate of the proportion of variance explained by the line effect in our generalised linear mixed models. This estimate was obtained with the “rptGaussian” function of the rptR package (Stoffel et al., 2017), with models including line, subline, individual and session as random effects, and line as a grouping factor.

#### Do personality traits differentiate the isogenic lines?

We compared the personality scores of the 24 lines, taking into account variation due to individual, subline and session effects. With the values of each personality score extracted from the PCA (see above), we first fitted a linear mixed-effects model with the “lmer” function of the lme4 package (Bates et al., 2015), with line as a fixed effect and individual, subline and session effects as random effects. We performed a Tukey all-pairs comparison on lines with the “glht” function of the multcomp package (Hothorn, Bretz, & Westfall, 2008). We graphically represented the distribution of each line along the two personality scores, for the same PCA as described above, estimated from individual values. We then used the “plot.PCA” function of the FactoMineR package to represent only mean point values for each line on the graph.

#### Are personality traits genetically correlated with fitness-related traits?

We investigated the genetic correlation between genetic traits, using the same linear mixed model as described in the introduction to this section. We first extracted linear predictors for each line and trait, with the “ranef” function of the lme4 package. We then used these values to calculate Spearman’s rank correlation coefficients. We estimated bootstrapped 95% confidence intervals, to assess significance of the Spearman’s rank correlation coefficients, and adjusted the *p*-values as described above.

## Results

### Are the measured behavioural traits repeatable?

Repeatability estimates for the seven behavioural traits ranged from 0.04 to 0.35 (Table 1). The repeatability estimates had confidence intervals excluding zero for all traits except for “time to first crossing of the border between the white and black areas” (Table 1). Only repeatable traits were considered in the subsequent analysis.

**Table 1.** Estimated repeatability (R) and 95% confidence intervals (between square brackets) for behavioural traits. Repeatable traits (R-value in bold type) were used to estimate personality scores.

| Personality trait category | Variable assessed | R [95% CI] |
| --- | --- | --- |
| **Activity** | Mean speed in area 4 | **0.35** [0.29; 0.40] |
|  | Activity rate in area 4 | **0.08** [0.01; 0.14] |
| **Boldness** | Change of speed in the border area (area 2) | **0.10** [0.04; 0.17] |
|  | Change of sinuosity in the border area (area 2) | **0.12** [0.04; 0.19] |
|  | Time to first crossing of the white/black border | 0.04 [0.00; 0.11] |
| **Exploration** | Sinuosity in area 1 | **0.24** [0.17; 0.30] |
|  | Area explored in area 1 | **0.18** [0.12; 0.24] |

### Do the measured traits identify individual behavioural syndromes?

All repeatable variables were correlated with at least one other variable (Table 2), indicating the existence of a behavioural syndrome. We combined these six variables into two personality scores based on the first two axes of a PCA, which accounted for 56.8% of the variance (Table 3). The first axis (personality score 1, PC1) was positively correlated with the “area explored in area 1” and inversely correlated with “sinuosity in area 1” and with the “change of sinuosity in the border area 2” (Table 3). Highly positive values of PC1 corresponded to a high exploration score (Figure 3). The second axis (personality score 2, PC2) and correlated mostly with “mean speed in area 4”, “activity rate in area 4” and the “change of speed in border area 2” (Table 3). High positive values of PC2 correspond to high activity scores (Figure 3).

**Table 2.** Phenotypic correlation between behavioural variables, with Spearman’s rank correlation coefficient Rho and 95 percent confidence intervals (between square brackets), based on a number of individual values from N = 977 to N = 1009. Correlation coefficients with confidence intervals excluding zero are shown in bold, and correlation coefficients remaining significantly different from zero after Benjamini and Hochberg correction are indicated with an asterisk. The personality trait category to which each variable belongs is indicated in brackets: activity (A), boldness (B) and exploration (E).

|  | (B) Change of speed in border area 2 | (A) Mean speed in area 4 | (A) Activity rate | (B) Change of sinuosity in border area 2 | (E) Sinuosity in area 1 |
| --- | --- | --- | --- | --- | --- |
| (A) Mean speed in area 4 | **0.31**<br>**[0.25; 0.37] \*** |  |  |  |  |
| (A) Activity rate in area 4 | **0.10**<br>**[0.04; 0.16] \*** | **0.38 \***<br>**[0.32; 0.43] \*** |  |  |  |
| (B) Change of sinuosity in border area 2 | **0.11**<br>**[0.05; 0.17] \*** | **0.07**<br>**[0.01; 0.14] \*** | **-0.12**<br>**[-0.18; -0.06] \*** |  |  |
| (E) Sinuosity in area 1 | **-0.07**<br>**[-0.14; -0.01] \*** | **0.13**<br>**[0.07; 0.19] \*** | **-0.16**<br>**[-0.22; -0.10] \*** | **0.38**<br>**[0.32; 0.44] \*** |  |
| (E) Area explored in area 1 | **0.11**<br>**[0.04; 0.16] \*** | 0.01<br>[-0.04; 0.08] | **0.29**<br>**[0.23; 0.34] \*** | **-0.28**<br>**[-0.34; -0.22] \*** | **-0.56**<br>**[-0.61; -0.52] \*** |

**Table 3.** Parameters from the first two principal components (PC1 and PC2) of the PCA for the behavioural variables measured. Component loadings represent the relationship between the principal components and the variables from which they are constructed. The personality trait category to which each variable belongs is indicated in brackets: activity (A), boldness (B) and exploration (E).

| Parameter | PC1 | PC2 |
| --- | --- | --- |
| Eigenvalue | 1.87 | 1.54 |
| Percentage of variance explained | 31.23 | 25.58 |
| Component loading |  |  |
| (A) Mean speed in area 4 | 0.16 | 0.84 |
| (A) Activity rate in area 4 | 0.43 | 0.60 |
| (B) Change of speed in area 2 | 0.19 | 0.51 |
| (B) Change of sinuosity in area 2 | -0.56 | 0.34 |
| (E) Area explored in area 1 | 0.81 | -0.09 |
| (E) Sinuosity in area 1 | -0.81 | 0.27 |

**Figure 3.**
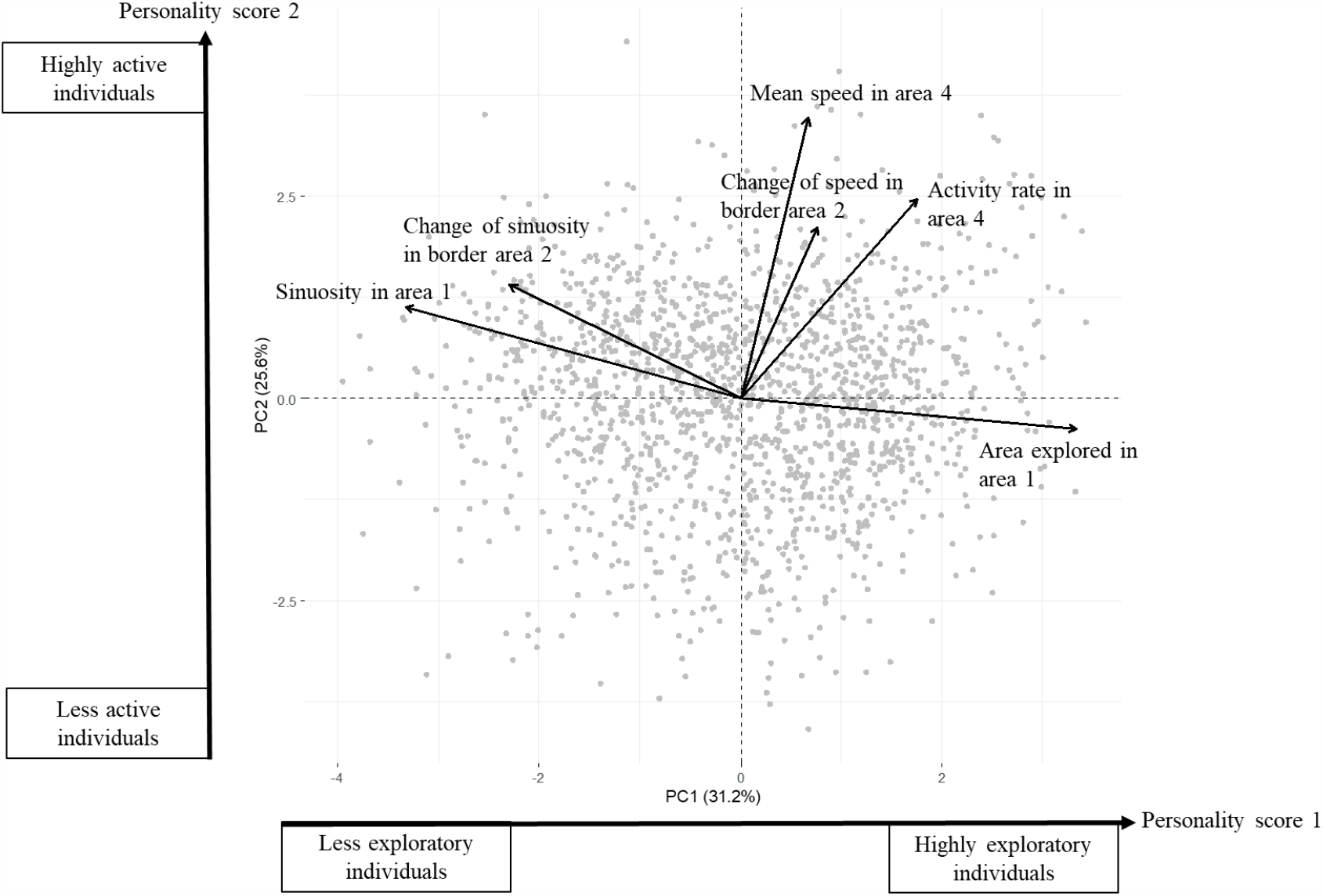
Graphical representation of the first two axes of the PCA on individual values (grey points) for repeatable behavioural traits (in black type).

### Are the measured traits correlated with fitness-related traits?

Active females (i.e. those with higher PC2 values) had significantly larger numbers of offspring and significantly longer tibias (Table 4). Higher rates of exploration (i.e. higher PC1 values) were not significantly correlated with any of the fitness-related traits measured. None of the behavioural variables or personality scores was significantly correlated with longevity (Table 4).

**Table 4.** Phenotypic correlation between behavioural traits (behavioural variables and personality scores) and other life history traits (with Spearman’s rank correlation coefficient Rho and 95% confidence intervals (between square brackets) calculated from 959 individual values). Correlation coefficients with confidence intervals excluding zero are shown in bold, and correlation coefficients that remained significantly different from zero after Benjamini and Hochberg correction are indicated with an asterisk. The personality trait category to which each variable belongs is indicated in brackets: activity (A), boldness (B) and exploration (E).

|  | Offspring number | Longevity | Tibia length |
| --- | --- | --- | --- |
| **Behavioural variables** |  |  |  |
| (A) Mean speed in area 4 | **0.20 [0.14; 0.26] \*** | -0.05 [-0.12; 0.03] | **0.19 [0.12; 0.25] \*** |
| (A) Activity rate in area 4 | -0.01 [-0.08; 0.06] | -0.06 [-0.13; 0.02] | -0.07 [-0.13; 0.00] |
| (B) Change of speed in border area 2 | **0.13 [0.06; 0.19] \*** | -0.08 [-0.15; 0.00] | **0.16 [0.10; 0.21] \*** |
| (B) Change of sinuosity in border area 2 | **0.11 [0.04; 0.17] \*** | 0.002 [-0.07; 0.08] | 0.05 [-0.01; 0.11] |
| (E) Area explored in area 1 | -0.05 [-0.11; 0.01] | -0.05 [-0.13; 0.02] | -0.02 [-0.09; 0.04] |
| (E) Sinuosity in area 1 | 0.01 [-0.05; 0.07] | 0.05 [-0.02; 0.13] | 0.05 [-0.01; 0.11] |
| **Personality scores** |  |  |  |
| Exploration score (PC1) | -0.01 [-0.07; 0.06] | -0.05 [-0.13; 0.03] | -0.03 [-0.09; 0.03] |
| Activity score (PC2) | **0.17 [0.10; 0.23] \*** | -0.01 [-0.10; 0.07] | **0.15 [0.09; 0.21] \*** |

### Are the measured traits heritable?

Broad-sense heritability estimates for behavioural traits and personality scores ranged from 0.01 to 0.11. Confidence intervals excluded zero for all traits linked to activity and exploration, whereas they included zero for the two traits linked to boldness (Table 5). Fitness-related traits (offspring number, tibia length and longevity) displayed broad-sense heritability ranging from 0.04 to 0.28, with all confidence intervals excluding zero (Table 5).

**Table 5.** Broad-sense heritability (H^2^) of traits measured with 95% confidence intervals (between square brackets). Heritability estimates are shown in bold if their 95% confidence interval did not include zero. The personality trait category to which each behavioural variable belongs is indicated in brackets: activity (A), boldness (B) and exploration (E). are considered to be significantly different (with a p-value <0.05).

| | $H^2$ [95% CI] |
| --- | --- |
| **Behavioural variables** |  |
| (A) Mean speed in area 4 | **0.11 [0.05; 0.18]** |
| (A) Activity rate in area 4 | 0.02 [0.00; 0.04] |
| (B) Change of speed in border area 2 | 0.01 [0.00; 0.03] |
| (B) Change of sinuosity in border area 2 | 0.01 [0.00; 0.03] |
| (E) Area explored in area 1 | **0.06 [0.02; 0.10]** |
| (E) Sinuosity in area 1 | **0.06 [0.02; 0.11]** |
| **Personality scores** |  |
| Exploration score (PC1) | **0.08 [0.03; 0.13]** |
| Activity score (PC2) | **0.05 [0.02; 0.10]** |
| **Fitness-related traits** |  |
| Offspring number | **0.12 [0.05; 0.19]** |
| Tibia length | **0.05 [0.01; 0.09]** |
| Longevity | **0.28 [0.14; 0.39]** |

### Do personality traits differentiate between lines?

We found significant differences in personality scores between lines (Figure 4.a and 4.b), and the 24 lines were distributed along the first two axes of the PCA (Figure 5). We were therefore able to distinguish between lines that were very active and exploratory (e.g., lines 3 and 12), and lines that were less active and exploratory (e.g., lines 14 and 21); we were also able to distinguish between lines that were very exploratory but not very active (e.g., lines 9 and 10) and lines that were active but not very exploratory (for example line 4).

**Figure 4.**
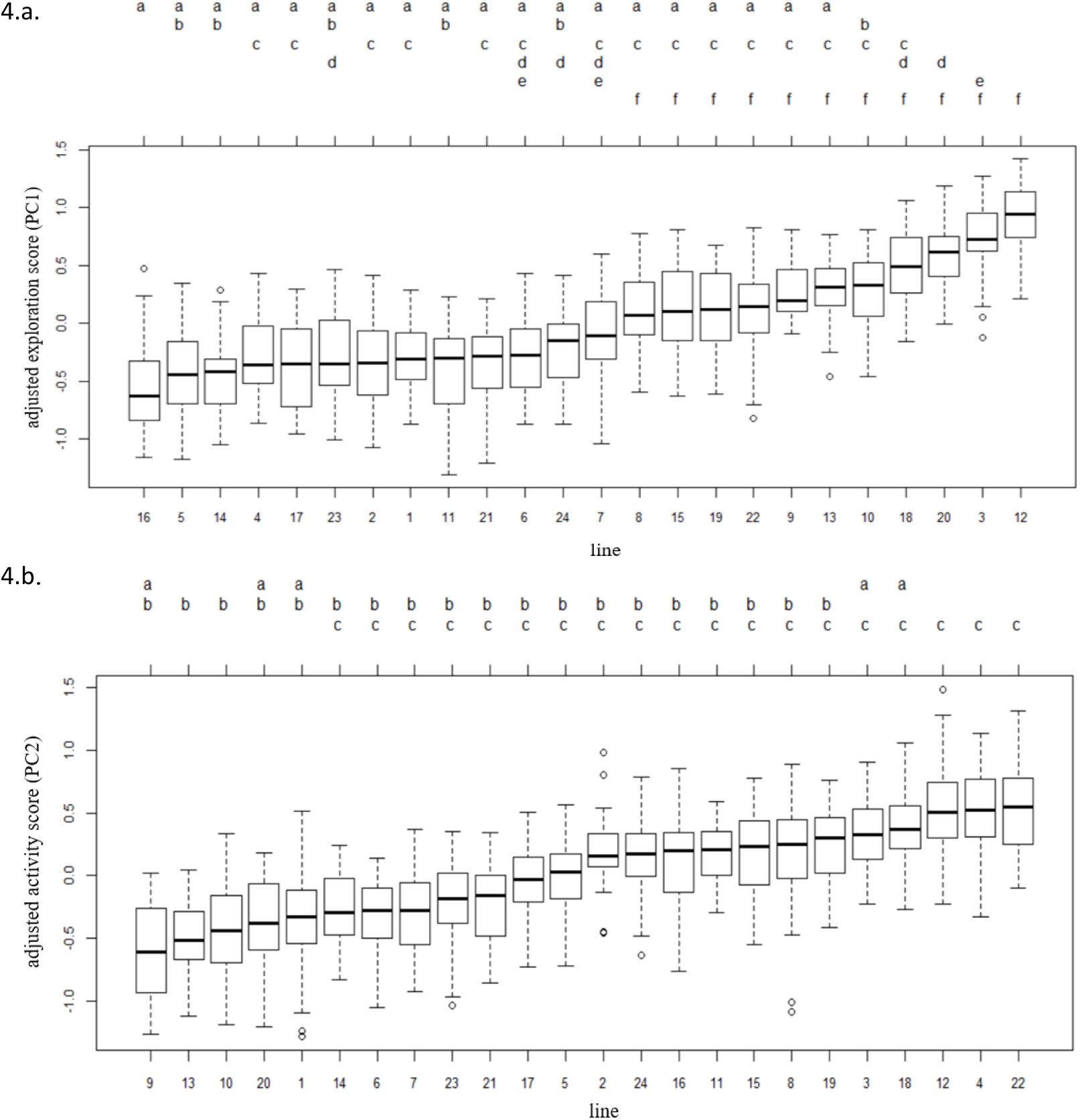
Boxplot of the adjusted values of personality score 1 (Fig. 4.a) and personality score 2 (Fig. 4.b) after the elimination of variation due to individual, subline and session effects, and compact letter display after Tukey all-pair comparisons. Two lines with no letters in common are considered to be significantly different (with a p-value <0.05).

**Figure 5.**
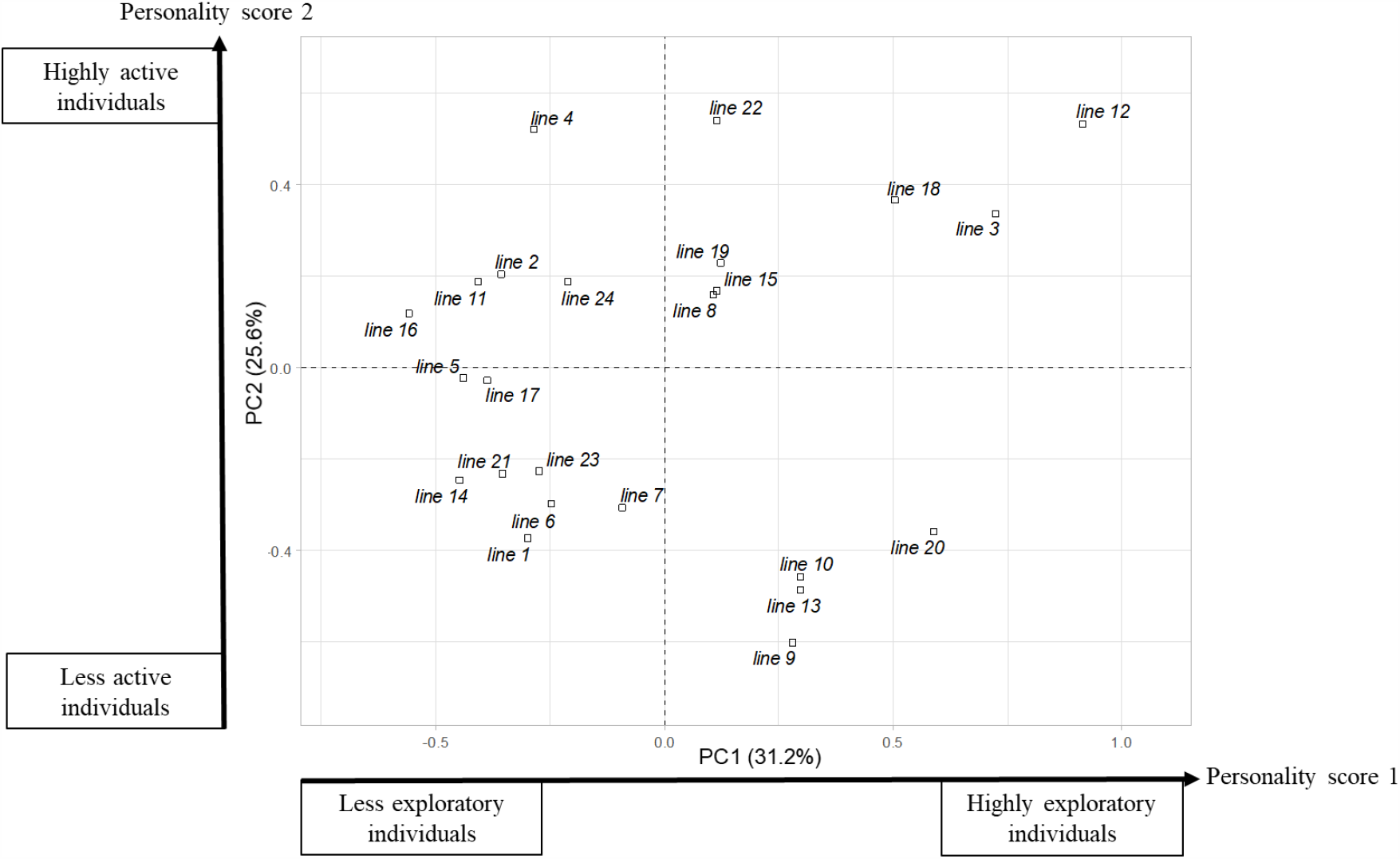
Distribution of the mean points for the 24 lines (centroids) along the first two axes of the PCA.

### Are personality traits genetically correlated with fitness-related traits?

The only genetic correlation between personality scores and fitness-related traits that remained significant after FDR correction was the positive correlation between exploration score (PC1) and offspring number (Table 6).

**Table 6.** Genetic correlation between personality and other life history traits (Spearman’s rank correlation coefficient, based on the trait estimates of 24 near-isogenic lines, with associated p-values in brackets). Correlation coefficients with confidence intervals excluding zero are shown in bold, and correlation coefficients that remained significantly different from zero after Benjamini and Hochberg correction are indicated with an asterisk. The personality trait category to which each behavioural variable belongs is indicated in brackets: activity (A), boldness (B) and exploration (E).

|  | Offspring number | Longevity | Tibia length |
| --- | --- | --- | --- |
| Behavioural variables |  |  |  |
| (A) Mean speed in area 4 | -0.16 [-0.54; 0.33] | -0.32 [-0.62; 0.06] | **0.51 [0.21; 0.70]** |
| (A) Activity rate in area 4 | **-0.53 [-0.78; -0.14]** | 0.15 [-0.28; 0.54] | -0.21 [-0.62; 0.28] |
| (B) Change of speed in border area 2 | -0.03 [-0.44; 0.37] | **-0.45 [-0.72; -0.10]** | 0.35 [-0.05; 0.63] |
| (B) Change of sinuosity in border area 2 | 0.29 [-0.11; 0.63] | 0.15 [-0.31; 0.56] | 0.19 [-0.20; 0.54] |
| (E) Area explored in area 1 | **-0.60 [-0.79; -0.26] \*** | 0.01 [-0.40; 0.43] | -0.28 [-0.65; 0.16] |
| (E) Sinuosity in area 1 | **0.63 [0.33; 0.82] \*** | 0.25 [-0.20; 0.62] | -0.03 [-0.43; 0.39] |
| Personality scores |  |  |  |
| Exploration score 1 (PC1) | **-0.64 [-0.83; -0.29] \*** | -0.09 [-0.52; 0.32] | -0.19 [-0.58; 0.24] |
| Activity score 2 (PC2) | -0.10 [-0.52; 0.39] | -0.22 [-0.66; 0.19] | **0.41 [0.01; 0.67]** |

## Discussion

We investigated whether animal personality could be used to develop or improve phenotyping methods for the BCA *Trichogramma evanescens*. We first developed an automated phenotyping method based on automated pathway analysis, providing a set of behavioural trait measures that proved repeatable over time and heritable (i.e. personality traits). We then identified differences in life history strategies between individuals as behavioural traits were correlated together and combined them into personality scores, which were correlated with other life history traits. Finally, we observed differences in personality scores between the 24 genotypes of *T. evanescens* and found a negative genetic correlation between exploration and fecundity.

### Evidence of personality traits in Trichogramma evanescens

Personality has never before been assessed in a species as small as *Trichogramma evanescens*. Based on other video-tracking studies in other species (Branson et al., 2009; Charalabidis, Dechaume-Moncharmont, Petit, & Bohan, 2017), we designed and developed a video-tracking approach measuring a large number of variables relating to the movements of *T. evanescens* individuals during their presence in the different areas (white, black and border areas) within an experimental arena. Here, we chose to work on seven variables that (i) could be calculated with methods commonly used in trajectory and movement studies (speeds, trajectory length and sinuosity estimates) (Mclean & Volponi, 2018) and (ii) we considered to be associated with some of the commonly defined personality traits defined by Réale et al. (2007): boldness, exploration and activity.

For each of the seven behavioural variables, we assessed repeatability, broad-sense heritability and phenotypic and genetic correlations between personality traits and between these traits and other life history traits, according to methods generally used in animal personality studies (Réale et al., 2007). For six of the seven behavioural variables, we observed significant repeatability (R) (values ranging from 0.10 to 0.35, Table 1). These six variables could therefore be considered as personality traits. The R values obtained were within the range of R values commonly observed for behavioural traits, although most were lower than the mean R value obtained for animal behaviour (0.35) (Bell et al., 2009). However, personality has rarely been studied in parasitoid insects, and a recent study on the parasitoid wasp *Venturia canescens* reported a similar R value for activity and a lower R value for exploration (about 0.10, whereas we obtained R values for exploration-related variables of 0.18 and 0.24(Gomes et al., 2019)).

The broad-sense heritability of the variables (ranging from 0.06 to 0.11, Table 5) was lower than the mean value for animal behaviour (0.24) in the meta-analysis by Dochtermann et al. (2019). Stirling et al. (2002) found no significant differences in heritability between behavioural and life-history traits in their meta-analysis, whereas we found that heritability values for personality traits were lower than heritability values of two classical fitness-related traits (offspring number and longevity) in *T. evanescens* (Table 5).

Behavioural traits could be grouped together into two continuums or behavioural syndromes (Denis Réale et al., 2007; Sih et al., 2004; Sih, Cote, Evans, Fogarty, & Pruitt, 2012): a continuum extending from individuals with low levels of exploratory behaviour to highly exploratory individuals, and a continuum extending from individuals with low levels of activity to highly active individuals (Figure 3). Bold (or shy) behaviour and active behaviour have been shown to be correlated with fecundity traits in several species (Biro & Stamps, 2008), but rarely in insects (Monceau et al., 2017). In this study, we found a weak but significant phenotypic correlation between behavioural traits, fecundity and body length, as shy or active females produced more offspring, and had longer tibias (Table 4). The positive correlation between activity (with the variable “mean speed”) and the length of tibia is quite intuitive, as it should be easier for individuals with longer tibia to cover larger distance. Moreover, bigger females would have more energy to spend for both offspring production and activity. However, although these positive correlations might have been expected, they are equivocal in the literature and seem to depend on the function of personality traits in a given species (Biro & Stamps, 2008; Gu, Hughes, & Dorn, 2006). We can note that the variable for shyness on which we found a phenotypic correlation with fecundity and tibia length is the “change of speed in border area 2”, which is also directly linked with speed abilities. Finally, an analysis of genetic correlations showed that the lines with the most exploratory individuals had the smallest numbers of offspring (Table 6). These correlations seem to be compatible with the pace-of-life syndrome (POLS) hypothesis, a currently debated hypothesis (Royauté, Berdal, Garrison, & Dochtermann, 2018), according to which, behavioural traits are related to morphological, physiological and other life-history traits (Réale et al. 2010).

### Potential of personality traits for use in genetic improvement of biocontrol agents

In this study, our aim was to evaluate the possibility of using personality traits as traits of interest in biological control, and of integrating these traits into genetic improvement programmes for the BCA *T. evanescens*. The six repeatable behavioural traits we measured were correlated with each other, and could be combined into two continuums. For each individual and continuum, we estimated a personality score corresponding to the position of the individual along the continuum, a common method in animal personality studies (Mazué et al., 2015; Monceau et al., 2017). We found that it was possible to capture a large proportion of the behavioural trait variance with two scores (36.2% of the total variance explained by personality score 1, and 26.4% explained by personality score 2). This finding highlights the utility of calculating a few synthetic indices (or scores), rather than measuring large numbers of variables, to obtain relevant information for BC. We therefore systematically present our results considering all the traits individually and summarized as two personality scores.

The relevance of the behavioural traits or synthetic scores to the context of BC was demonstrated by the phenotypic correlations between these traits and scores and the traits classically measured in BC (fecundity, longevity and body length) (Hopper et al., 1993; Prezotti et al., 2004; Roitberg et al., 2001; Smith, 1996). In this study, active females (i.e. with high values for “mean speed in area 4” and “personality score 2”) produced more offspring and had longer tibias (Table 4). By contrast, we found that bold females (i.e. with low values for “change of speed in border area 2” and “change of sinuosity in border area 2”) produced a small number of offspring (Table 4). In several species, activity and boldness behaviours have been shown to be correlated with traits of ecological importance, such as dispersal (Sih et al., 2004), which is also a trait linked to field efficiency in BC (Fournier & Boivin, 2000). Our results indicate that active females produce more offspring, which is predictive of a high degree of efficiency in rearing conditions and, in the case of parasitoids, in the field. Note, however, that we did not assess survival or body condition in the offspring. The same females also displayed shyer behaviour. The impact of a shy behaviour on an individual’s field efficiency would depend on the agrosystem conditions. Indeed, in the presence of high densities of predators intraguild predation may occur (Bennett, Gillespie, Shipp, & VanLaerhoven, 2009; Dumont et al., 2018). In this scenario, shy parasitoid individuals (i.e. the intraguild prey) might be less predated as they might be less willing to take risks, compared to bold individuals. However, in situations where intraguild predation is not a challenge, bolder individuals, more willing to take risks, could be faster in finding resources (i.e. egg patches in the case of *Trichogramma* species). Therefore, further studies are required to assess the full ecological relevance of the lines we studied in BC. The relevance of the variables measured will be confirmed only if they are shown to be correlated with BC performance in industrial and field and/or greenhouse conditions.

Most of our data analyses aimed to evaluate the added value of the measured behavioural traits for genetic improvement strategies, breeding programs. We found that personality scores differ among isogenic lines (Figure 4.a. and Figure 4.b.) and that these differences highlight contrasted behaviours, as evidenced by their distribution along the two personality scores in Figure 5. This may make it possible to differentiate between these behaviours and to select for them, should they prove relevant in terms of BC efficiency. We also observed a negative genetic correlation between the personality score relating to exploration and offspring production. It will probably be important to take this trade-off into account in BC, as it may oppose performance in rearing and performance in the field. Indeed, as for activity and boldness, exploration behaviours are also correlated with traits linked to field efficiency in BC, such as dispersal (Fournier & Boivin, 2000; Sih et al., 2004).

Given these results, and the ease with which all the traits can be assessed and personality scores obtained through short (90 seconds) automated video-tracking measurements, the new method described here may provide useful criteria for the selection of candidate BCA taxa (populations, strains, sibling species, etc.) or for quality control purposes. However, the high level of intra-isogenic line variability observed (Figure 4.a and Figure 4.b), accounting for the relatively low broad-sense heritability of the traits and scores (between 0.01 and 0.11; Table 5), constrains the use of this method, as it may be necessary to phenotype large numbers of individuals for reliable comparisons between taxa or reared populations. The low heritability also constitutes an obstacle to the implementation of ambitious experimental evolution programmes. Oriented experimental evolution may be fastidious for traits displaying such a high degree of environmentally induced variability. As a comparison, breeding programmes for livestock animals generally make use of traits with higher heritability. Heritability values for morphological, physiological, behavioural or other traits linked to fitness and considered in these breeding programmes generally range from 0.17 to 0.70 in sheep, pigs, cows and fish (Juengel et al., 2019; Kavlak & Uimari, 2019; Moretti, de Rezende, Biffani, & Bozzi, 2018; Vargas Jurado, Leymaster, Kuehn, & Lewis, 2016). However, in order to select traits with low heritability values, the method of genomic selection is already used for livestock animals (e.g. Hayes, Bowman, Chamberlain, & Goddard, 2009). This method is based on the phenotyping and genotyping of a high number of individuals in order to establish a statistical equation between the genotype and the phenotype. Based on this equation, it is then possible to predict the phenotype of an individual, knowing only its genotype (Hayes et al., 2009). This method has never been applied to BCA, but has been recently suggested as a promising application to BCA selection (Leung et al., 2020), and could help considering behavioural – and personality – traits in BCA selection programs.

## Conclusion

In conclusion, the use of methods and concepts of animal personality to develop phenotyping methods and associated data analyses for BC led to the rapid phenotyping of traits rarely used in BC that were repeatable, heritable and correlated with fitness-related traits. Our results also provide support to investigate the interest of animal personality in other BCA species (parasitoids or predators). However, it will be possible to consider the actual potential of these traits and of the phenotyping method satisfactory only after investigating the relationships between the laboratory-measured traits and BC performance indices in real BC situations, in industrial production settings or in field releases. This first study has driven the launch of large-scale field experiments, which are currently underway and aim to generate field-release performance indices.

## Data accessibility

Data tables and code needed to re-do the analyses and figures are available online on Zenodo (http://doi.org/10.5281/zenodo.4058218).

## Acknowledgements

We would like to thank all the members of staff at Institut Sophia Agrobiotech who helped us when we were overwhelmed by the sheer numbers of *Trichogramma* individuals in our experimental tubes, including Léa Tregoat-Bertrand in particular. Special thanks to Maxime Dahirel for helpful advice for statistical analysis, and to Morgane Bequet-Rennes and Simon Vaïsse, who contributed to this study by running previous experiments that led to our decision to design this new phenotyping assay. We thank the biocontrol company BIOLINE, in particular Paloma Martinez and Benoît Franck, and the INRAE biological resource centre “Egg-Parasitoids Collection” (CRB EP-Coll, Sophia Antipolis) for providing field-sampled populations for the establishment of some of the isogenic lines used in this study. We thank the Microscopy Platform of the Sophia Agrobiotech Institute, INRA, UNS, CNRS, UMR 1355-7254, INRA PACA, Sophia Antipolis for providing access to instruments and technical advice. This work was funded by the the French “Agence Nationale de la Recherche” under the “Investissements d’Avenir UCAJEDI” project with reference n ° ANR-15-IDEX-01. Silène Lartigue received financial support from the French Ministry in charge of Agriculture. Version 4 of this preprint has been peer-reviewed and recommended by Peer Community In Ecology (https://doi.org/10.24072/pci.ecology.100069). We thank Marta Montserrat, Bart A. Pannebakker, François Dumont, Joshua Patrick Byrne and Ana Pimenta Goncalves Pereira for their thoughtful comments.

## Appendix

The initial populations were established from individuals sampled in different parts of France, from 2010 to 2016, by the INRAE “Egg-Parasitoids Collection” (CRB EP-Coll, Sophia Antipolis) biological resource centre or the company Bioline AgroSciences Ltd. The genetic and phenotypic differences between these populations were not known. As a means of obtaining populations with low levels of genetic variability, we established 14 lines from the initial populations, by brother × sister crossing over 15 generations (Figure 6.a.). As a means of obtaining lines with mixed genetic backgrounds, we established 10 lines from a mixture of three initial field-sampled populations. We achieved this by mating one virgin female from a first population with one virgin male from a second population. We then selected a virgin female from their offspring and mated it with a virgin male from a third population (Figure 6.b.). Over the next 13 generations, we performed inbred crosses, as explained above. The experiments described in this study were performed with these 24 lines.

**Figure 6.**
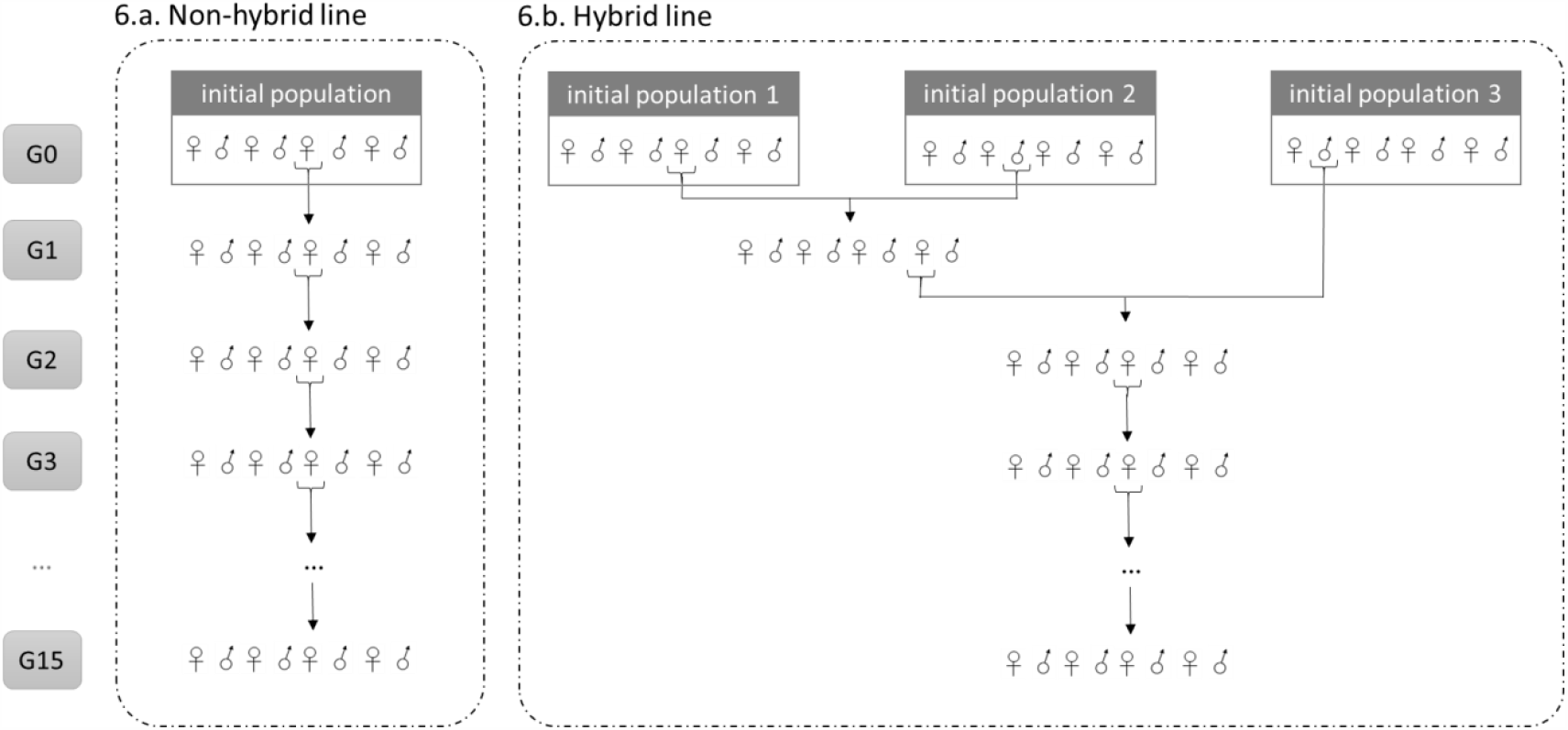
Crossing design for non-hybrid (Fig. 6.a) and hybrid (Fig. 6.a) lines, over 15 generations (G1 to G15).

